# The aptabot: an inducibly affinity-switching, minimally invasive *in vivo* contrast agent

**DOI:** 10.1101/043950

**Authors:** Elleard Felix Webster Heffern, Jason Fuller, Russell W. Hanson

## Abstract

The ambitions of current neuroscience—understanding neurological disease progression and mapping the connectome—demonstrate a need for safe *in vivo* tools for creating intricate maps of brain circuitry. Present *in vivo* contrast agents are often limited by their specificity, uptake, resolvability, and/or clearance.

We describe an aptamer-functionalized sensor for high-resolution imaging that can switch imaging targets by an induced multi-stage aptamer reaction. Included are synthetic methods as well as calculations of sensor efficacy based on known kinetics. Calculations show that 10 distinct targets may be imaged in a living brain at the submicron scale within 42 hours.

## Introduction

In order to map fine details of brain circuitry, agents must be developed that both bind specific targets of interest and provide highly resolvable contrast. MRI contrast agents have long been used to image brain activity and structure, and in recent years, specifically targeted PET, MRI, and radio imaging agents have been developed for expressed cell surface markers (Xue 2009). Current whole brain contrast agents are often constrained either by radiolabels that provide inadequate resolution to identify individual receptors and emit radiation in their environment, or metal nanoparticles with inefficient clearance or degradation that impedes imaging of subsequent targets. A multi-functionalized contrast agent that attaches to successive targets can provide the key to more intricate brain maps.

To overcome limitations in brain imaging, we propose creating smarter contrast agents by combining current nanotechnological principles. We call these agents "aptabots," based on their composition as aptamers and their mechanistic capacities at the nanoscale. Aptabots meet several criteria: A) specified target binding; B) traversal of the blood-brain barrier (BBB); C) highly resolvable contrast provision; and D) capacity for easy clearance. By achieving each criterion, aptabots can map the relative locations of multiple targets in single brains in great detail, without harming test subjects.

We propose affixing nanoparticles to our aptabots to confer agent stability and enable the use of high-resolution, bone-penetrating imaging techniques such as MRI or X-ray CT to visualize agent distribution. Further, we propose employing aptamers (specifically, stabler L-DNA spiegelmers) targeted to neuroproteins of interest coupled with simple machines driven by ssDNA strand displacement reactions can provide molecularly inducible affinity gain/loss. The choice of aptamers over chemical ligands as affinity-providing components eases the addition of ssDNA displacement machinery and makes inhibition as well as subsequent activation of affinity possible.

Aptabots can be constructed with existing conjugation methods and use simple DNA nanomachines with well-studied kinetics to provide contrast for specific targets and also allow for binding structures to gain and lose affinity. Furthermore, aptabots can incorporate aptamers that facilitate receptor-mediated transmission through the blood-brain barrier. The use of aptabots should dramatically improve *in vivo* brain imaging capacities.

**Inset 1.**
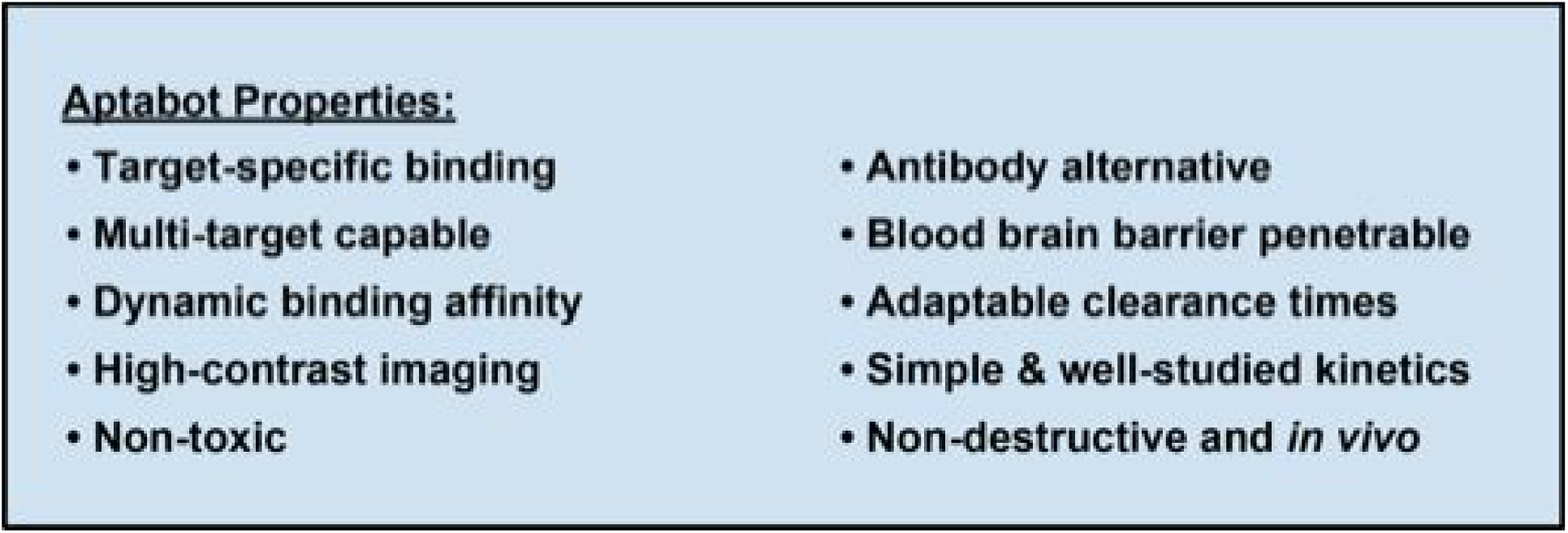

### Nanoparticles as contrast agents for high-resolution imaging of the *in vivo* brain

Several options exist to provide contrast for varied tissue structures. Nanoparticles are small particles ranging in size between 10 and 200 nm that are used both for contrast provision and as platforms to stabilize otherwise biodegradable molecules. Metallic nanoparticles can provide contrast for sub-micron resolution, skull-penetrating imaging methods including X-ray CT (if gold or silver) and MRI (if superparamagnetic iron oxide or gadolinium) (Lee N 2013, Shokrollahi 2013). These particles can be highly functionalized via known conjugation methods both for targeting and for other desirable capacities (Lee JH 2013, Montenegro 2013, Jiao 2011, Cheng 2013). Coating the surface of a metallic nanoparticle or adjusting its surface charge averts toxicity problems (Nam 2013). Quantum dots and fluorophores are smaller than metallic nanoparticles and can be highly functionalized as well, but require optical methods that fail to penetrate the skull (Geszke-Moritz 2013, Cassette 2013, Small 2014). Radiolabels can be used for imaging via PET or for SPECT, both of which do penetrate the skull, but these methods develop lower-resolution images than necessary for mapping the connectome (Zhu 2014, Yamamoto 2013, Rahmim 2014, Beekman 2005). Thus, we propose a method for specific *in vivo* imaging to resolve neural anatomy at the cellular and sub-cellular level using engineered nano-scale contrast agents.

### The choice of aptamers to provide sensing affinity to nanoparticles

Aptamers are single-stranded DNA or RNA molecules that bind with high affinity and specificity and are capable of conjugation. Due to the propensity of DNA and RNA to form helices and loops, they can form a wide variety of shapes, and thus, aptamers can be developed to bind selectively to a broad range of targets of interest. Aptamers offer a similar range of specificity to synthetic chemical ligands, antibody proteins, or small peptides (Stoltenburg 2007, Jing 2010) and multiple methods have been established for conjugating aptamers to nanoparticles (Lee JH 2013).

Aptamers have myriad functional advantages over other affinity-providing molecules. Aptamers are capable of greater specificity and affinity than other binding molecules and can easily be modified to offer heightened structural specificity (Javasena 1999, Jing 2010, Ferreira et al 2008, Burbulis et al 2008). Aptamers are composed of single, self-refolding chains that low redox potential, allowing for tolerance to and recovery from traumatic pH or temperature shifts, allowing aptamers, unlike antibodies, to be transported without special cooling requirements.

The development of aptamers with affinity for custom in vivo targets is considerably more straight forward than antibody production, as aptamers are produced via chemical synthesis rather than through the use of bacteria culture and/or animals. Application of aptamers is less prone to elicit an immune response that interferes with binding processes than is the application of antibodies. Furthermore, attachment of an aptamer confers less volume to an imaging agent that might interfere with agent migration than does attachment of an antibody (McCauley et al 2003, Cao et al 2009, De Rosa and La Rotonda 2009, Ferreira et al 2009, Yan and Levy 2009). Most interestingly, because aptamers are composed of nucleic acids, they allow for conjugation via methods that can be exploited selectively to release conjugated nanoparticles from their targets.

Varied methods have been developed effectively to conjugate aptamers to nanoparticles. Different conjugation chemistries include streptavadin-biotin binding, thiol substitution reactions, and hybridization of extended DNA sequences concatenated to aptamers to so-called capture oligonucleotides that have been functionalized onto nanoparticle surfaces. Several of these methods are illustrated in Figure 1.

**Figure 1.**
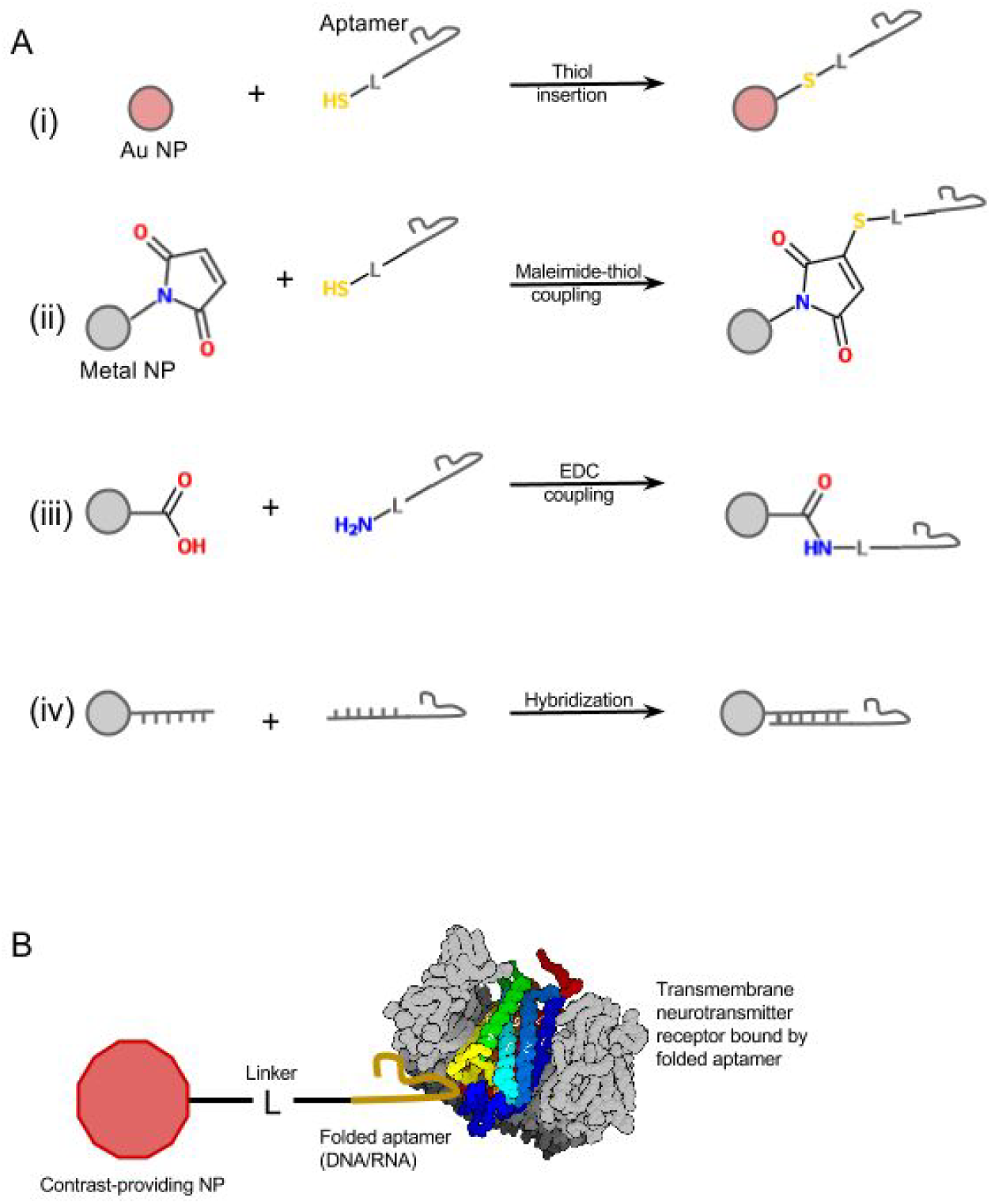
Nanoparticle-aptamer conjugation methods, and application of conjugates. **A.** Conjugation methods include i) thiol insertion on gold nanoparticles or ii) maleimide-thiol coupling on maleimide-tagged metal nanoparticles, with thiol-tagged aptamers, iii) EDC coupling on carboxyl-tagged metal nanoparticles with amino-tagged aptamers, and iv) hybridization of ONT-functionalized nanoparticles to aptamers with extended sequences. **B.** Nanoparticle-aptamer conjugates bind to targeted neurotransmitter receptors *in vivo* and provide imaging contrast.

Because they are nucleic acids, aptamers, are targets of bloodborne endonucleases; however, this can be addressed in several ways. Modifications may be made to developed aptamers, generally including fluoro or amino additions at the 2’ sugar centers of the DNA chains, enhance their resistance to these enzymes without strongly affecting folding tendencies or binding affinities (Javasena 1999). Alternatively, L-nucleic acid chains known as spiegelmers may be created via a similar process to SELEX to confer even greater endonuclease resistance.

Since the technology in this paper uses similarly enantiomeric free nucleic acid chains (“substitute strands”) to un-inhibit aptamer folding and subsequently to release aptamers from agents, the use of spiegelmers may in fact be necessary for the technologies illustrated in this paper, since these substitute strands will similarly have greater endonuclease resistance if they are L-enantiomers. If D-DNA aptamers with 2’ sugar modifications are used, then substitute strands may be stabilized via concatenated extensions and inexpensive polyorganic nanoparticle platforms.

### Time-saving advantages of inducible affinity switching

Imaging and mapping single targets *in vivo* is an impressive feat; elucidating intricate maps of brain circuitry *in vivo*, however, requires the imaging and mapping of multiple targets in single organisms. This demands either eliminating contrast agents from imaging targets once they are imaged, so that subsequent targets can be imaged with similar methods and minimal background, or subtracting prior images from new images once contrast agents with new affinities are added to the system. But since contrast is not linearly proportional to agent concentration, subtraction is rough and inaccurate.

Pharmacokinetic clearance times for stealthy metal nanoparticles indicate that the half-lives for concentrations of target-bound metal nanoparticle contrast agents may range from hours to weeks. For highly sensitive contrast agents, binding potential, the ratio between the concentration of ligands bound to targets and the total concentration of total ligands, ideally exceeds 1. In instances of low target occupancy, binding potential can be approximated to equal to the concentration of total targets divided by the dissociation constant. Thus, if a free nanoparticle has a blood concentration half-life of *n*, the concentration of targeted nanoparticles bound to targets may be given by the equation (see **Supplementary Mathematics**):

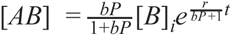

where [*AB*] is the concentration of agents bound to targets; [*B*] is the concentration of unbound agents, [*B*]_*i*_ is the initial concentration of unbound agents; *bP* is the binding potential; *r* is the product of *n* and the natural logarithm of ½; and *t* is time in days. Calculated from the above equation, the half-life, *T*_½_, of [*AE*] is given by the equation below:

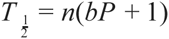

From these equations we can see that clearance rates for aptamer-conjugated agents become problematic when imaging subsequent targets. For instance, positing 30 nanomolar brain target concentration, and dissociation constants for protein-bound aptamers that range from low micromolar to high picomolar levels (Jing 2010), binding potential for targets will range from 1 to 300. According to 2D regressions, the blood concentration half-life of a stable untargeted ~20nm unPEGylated gold nanoparticle is approximately 4 hours, and this number increases when employing PEGylation or smaller particles (Dreaden 2012). Thus, if the half-life in the brain of a PEGylated, untargeted 20 nm gold nanoparticle with BBB transversal affinity equals 4 hours, then the half-life of the targeted version of this agent ranges from a minimum of 8 hours to 12 days. Enabling agents to lose affinity for their targets upon induction by non-endogenous small molecules helps overcome this problem by reducing half-lives from *n*(*bP* + 1)back to *n*. Thus when using a releasable agent, a brain clearance delay only of 4.32 multiplied by *n* is needed to ensure that 95% of these agents are cleared after agents inducibly lose affinity for their targets (0.5^4.32^=0.05). This means allowing approximately 17 hours for a small Au nanoparticle to clear.

Imaging multiple targets in a single brain at even higher rates could be achieved through molecular induction. An agent’s current target could be released and a new and different ligand could be exposed. Waiting for such an agent to clear would be unnecessary.

### Strand displacement reactions to invoke affinity gain and loss

Affinity-switching functions of aptabots are driven by toehold-mediated ssDNA strand displacement reactions. In a toehold-mediated strand displacement reaction, a removable ssDNA strand begins hybridized to a fully complementary base ssDNA strand that has a sequence that extends beyond the complementary region called a toehold (Figure 2**, left**). A substitute ssDNA strand with complete complementarity to the base strand including the toehold region is added to solution. The substitute strand first binds to the toehold sequence of the base strand, then progressively displaces the removable (incomplete complement) strand through a sequence of reversible single nucleotide-denaturation and single nucleotide-annealing reactions. (Figure 2**, middle**). Once the substitute and base strands fully duplex, no toehold exists for removable strands easily to re-initiate binding with base strand-substitute strand duplexes (Figure 2**, right**).

**Figure 2.**
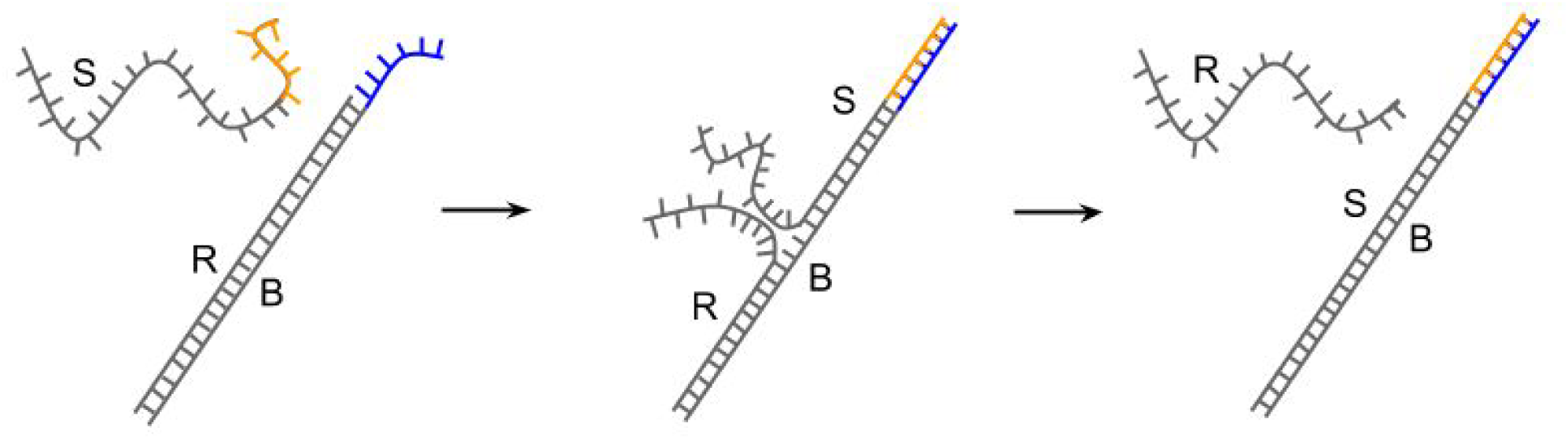
Toehold-mediated strand displacement reaction. R operates as the removable strand; S operates as the substitute strand; and B operates as the base strand.

Equilibrium constants for strand displacement reactions are affected by both toehold length and toehold sequence composition. The Gibbs free energy change of the reaction

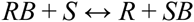

where *B* is the base strand, *S* is the substitute strand, and *R* is the removable strand can be calculated using the nearest-neighbor model of nucleic acid hybridization thermodynamics with values observed for sodium concentrations close to those in brain extracellular fluid. It falls within a range of approximately -1.99 kcal\**m* to -0.89 kcal\**m* per mole of reactants, where *m* is toehold length (Vologodskii 1984, SantaLucia 1998). Equilibrium constants based on these values range between 4.23^*m*^ to 25.2^*m*^, depending on sequence specifics. These parameters are significant as they directly affect the ratio of agents that are in an appropriate state after multiple strand displacement reactions.

Optimal pre-hybridized toehold sequence are composed either entirely of guanine or entirely of cytosine residues (post-hybridized, G…G bound to C…C). This composition nearly maximizes the magnitude of Gibbs free energy change of strand displacement (-1,64*m* kcal/mol) and hence the equilibrium constant (14.3^*m*^), while preventing hairpin formation in the pre-hybridized toehold. Pre-hybridized toehold hairpins are undesirable as they inhibit substitute strand-base strand binding. Table 1 considers the range of the ratio of agents on which every strand displacement reaction occurs to total agents, in the situation where 8 strand displacement reactions are sequentially induced, with respect to toehold length.

The kinetics of toehold-mediated strand displacement reactions are well-studied. The maximum rate of strand exchange shows an exponential dependence on toehold length, and hence the time for a certain proportion of strands to exchange, given equal starting concentrations of base strand-removable strand duplexes and substitute strands in solution, is exponentially dependent on the opposite of toehold length (Yurke 2003). These kinetics become important when choosing appropriate toehold lengths to ensure that strand displacement driven affinity switching saves a relevant amount of time in the imaging process. Generally, toehold length greater than or equal to 6 nucleosides ensures that each strand displacement reaction can proceed beyond 95% completion in 15 minutes. The use of this toehold length is referenced in Figures 5 and 6.

In our smart contrast agent design, toehold-mediated strand displacement can invoke either gain or loss of affinity. With respect to affinity gain, a set of strand displacement reactions removing “protector” ssDNA strands, exposing a ssDNA aptamer, allowing it to fold. Similarly, a ssDNA “attachment” strands that have been concatenated to the aptamer-folding strand, can be displaced by single strand displacement reactions to initiates affinity loss.

**Table 1.** Predicted range of the Gibbs free energy change of a single strand displacement reaction RB + S <-> R + SB, and range of the equilibrium constant, based on toehold length. Numbers in parentheses denote values calculated for when sequence of the toehold and the 5’ preceding nucleotide on the base strand consists either entirely of guanine residues or entirely of cytosine residues. *For all reactions or reaction sequences, a stoichiometric quantity (equimolar to starting RB concentration) of each reactant is added.

| Toehold length, m<br>(number of nucleotides) | $\Delta G$ range of each strand displacement reaction,<br>$RB + S \leftrightarrow R + SB$<br>(kcal/mol)<br>*The exact value of $\Delta G$ depends on the toehold sequence and the 5' preceding nucleotide.* | K range of each strand displacement reaction<br>*The exact value of K depends on the toehold sequence and the 5' preceding nucleotide.* | Ratio range of removable strand concentration ([R]) to removable strand-base strand duplex concentration ([RB]), at equilibrium*<br>*The exact value of this ratio depends on the toehold sequence and the 5' preceding nucleotide.* | Ratio range of removable strand concentration [R] to total concentration of removable strands ([R] + [RB]), at equilibrium*<br>*The exact value of this ratio depends on the toehold sequence and the 5' preceding nucleotide.* | Ratio range of agents where all strand displacement reactions have occurred, to total contrast agents, following induction of 8 strand displacement reactions*<br>*The exact value of this ratio depends on the toehold sequence and the 5' preceding nucleotides.* |
| --- | --- | --- | --- | --- | --- |
| 0 | 0 | 1 | 1 | 0.5 | 0.00391 |
| 1 | -0.89 to -1.99<br>(-1.64) | 4.23 to 25.2<br>(14.3) | 2.06 to 5.02<br>(3.78) | 0.673 to 0.884<br>(0.79) | 0.042 to 0.224<br>(0.153) |
| 2 | -1.78 to -3.98<br>(-3.28) | 17.9 to 635<br>(204) | 4.23 to 25.2<br>(14.3) | 0.809 to 0.962<br>(0.935) | 0.183 to 0.732<br>(0.582) |
| 3 | -2.67 to -5.97<br>(-4.92) | 75.7 to 16000<br>(2920) | 8.71 to 127<br>(54.1) | 0.897 to 0.992<br>(0.982) | 0.419 to 0.939<br>(0.864) |
| 4 | -3.56 to -7.96<br>(-6.56) | 320 to 403000<br>(41800) | 17.9 to 635<br>(204) | 0.947 to 0.998<br>(0.995) | 0.647 to 0.987<br>(0.962) |
| 5 | -4.45 to -9.95<br>(-8.20) | 1350 to<br>10200000<br>(598000) | 36.9 to 3190<br>(771) | 0.974 to 1.00<br>(0.999) | 0.807 to 0.997<br>(0.990) |

### Blood-brain barrier traversal

A challenge, both to effectively targeting brain protein with a contrast agent and to inducing strand displacement on an agent within the brain, is crossing the blood brain barrier. For blood-bound particles with minimal lipid solubility and/or large masses, the optimal method for blood-brain-barrier (BBB) penetration is to utilize receptor-mediated transcytosis (RMT), the same mechanism used to transport insulin into the brain (Pardridge 2005).

One option for RMT delivery is to package agents or substitute strands within biodegradable nanoparticle coats, as if they were drugs or genes (Panyam 2012, Pinzón-Daza 2013). However, ensuring coat degradation near targets within the brain is challenging. Alternatively, RMT-targeted antibodies, such as that for transferrin receptor (TfR), have been shown to facilitate contrast agent delivery across the blood-brain barrier of otherwise non-functional drug-bearing agents, and dual targeting with this method (where agents are also functionalized to bind to specific imaging targets, as would be the case with aptabots) has been shown to substantially enhance MRI performance of contrast-providing nanoprobes for intracranial blastomas (Gulati 2013, Ni 2014). However, while antibodies can be utilized as general RMT agents, they are not likely to work with substitute strand displacement due to the steric interference they will surely cause.

The optimal option for RMT delivery both of agents and of substitute strands appears to be the use of an aptamer that monovalently targets RMT mechanisms (Cheng 2013). This aptamer can be conjugated to any nanoparticle agent for dual-targeting, as would an antibody. And unlike an antibody, it also can be concatenated to any substitute strand at the non-toehold-containing terminus. The toehold-containing terminus of the substitute strand remains free and relatively sterically uninhibited to initiate strand displacement upon reaching its target. Further, although concatenating the RMT-targeting aptamer to the substitute strand is simple and inexpensive, if steric interference of the RMT-targeting aptamer inhibits strand displacement (and this can be tested on its own *in vitro* concurrently with testing effective BBB transport of the concatenation *in vivo*), then liposome packaging may be necessary as an alternative for substitute strand delivery.

Regarding agents themselves, additional steps may be taken to ensure appropriate nanoparticle access to the brain. Particularly, coating nanoparticles or adjusting surface charge prior to conjugation allows for stealthy evasion of immune mechanisms. Methods for coating various nanoparticles are fairly well-studied: one group demonstrated functional methods for coating gold nanoparticles in PEG without the aid of an additional polymeric dendrimer coat (Niidome 2006). On the other hand, adjusting surface charge is a less expensive and equally well-established process shown not to preclude subsequent aptamer conjugation (Kim 2010). Finally, it is important to consider that even RMT-dual-targeted nanoparticle complexes may show intolerably low levels of BBB penetration. Should such a circumstance arise, nanoparticle modification towards this end may be swapped for more aggressive methods to enhance general BBB penetration of large plasma-bound particles, such as mannitol injection (Wang 2007), or ultrasound permeabilization (Arvanitis 2012).

### Simple aptabots with affinity loss capacity

In Figure 3A, the simplest and perhaps most plausible model for the aptabot is illustrated: a contrast-providing nanoparticle is conjugated to an attachment strand of ssDNA hybridized to a link strand of ssDNA, which is concatenated to a folded aptamer. The attachment strand has a toehold region at the aptamer end. The link and attachment strands begin hybridized, linking the folded aptamer and any bound targets to the contrast-providing nanoparticle. Substitute strands with full complementarity to attachment strands are added to solution. This addition initiates toehold-mediated displacement of the link strand, freeing the contrast-providing nanoparticle to migrate from the target for systemic clearance (Figure 3C). The link strand and folded aptamer remain bound to their target without providing contrast. Future imaging will show only the distribution of new contrast agent added after strand-displacement.

**Figure 3.**
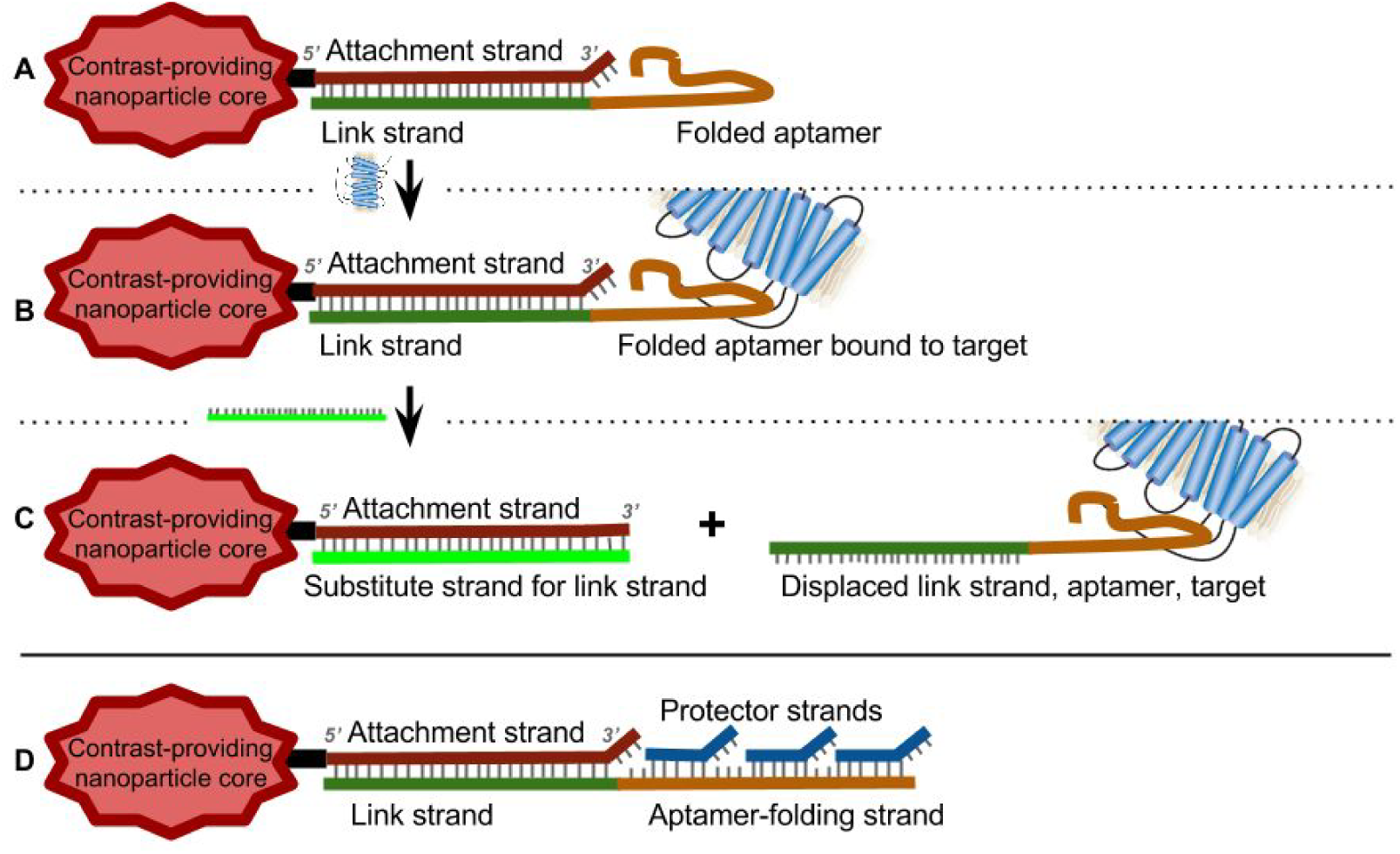
Simple aptabots with single affinity-changing arms. For the purpose of clarity, substitute strands in the figures are illustrated without concatenations to RMT-targeting aptamers. **A.** An aptabot with an arm capable of losing receptor affinity. A folded aptamer targeted to a transmembrane protein is concatenated at its to a link strand on its 3’ end; this link strand is hybridized to an attachment strand, which is conjugated to a core nanoparticle at its 5’ head and has a toehold region at its 3’ tail. **B.** Aptabot providing contrast for its target, due to aptamer-target binding after aptabot and target are mixed in solution. C. Aptabot after its arm loses receptor affinity, following addition to solution of a substitute strand fully complementary to the attachment strand, and subsequent displacement of the link strand by the substitute strand as the new binding partner to the attachment strand. The link strand, aptamer, and target protein are displaced from the complex, so the aptabot is free for clearance. **D.** An aptabot with a slightly more complex arm that begins without any target affinity. Protector strands, each with a toehold region, shield the aptamer strand and prevent it from folding; upon addition of strands with full complementarity to the protector strands, the affinity strand is displaced from the protector strands and folds into an aptamer, giving a conformation akin to that shown in **A**.

Contrast agents have been synthesized where the attachment strand, referred to as a “capture oligonucleotide”, has been functionalized onto gold nanoparticles for capture of ssDNA strand with single termini folded into aptamers; prostate cancer cells have adequately been targeted and imaged with contrast agents synthesized this way (Kim 2010). Constructing a simple nanobot as illustrated in **Figure 3A** requires only a minor adjustment of the synthetic procedure—extending the capture oligonucleotide to include a toehold stretch, prior to conjugation and hybridization.

While this paper proceeds to discuss higher-functioning aptabots with multi-stage capacities, the two-stage aptabot shown in **Figure 3A**, together with multifunctionalization for BBB penetration, provides perhaps the best combination of value and ease-of-construction. It requires adequate attachment only of two arms to the nanoparticle - that targeting RMT, and that targeting the imaging target - and addresses the problem of systemic clearance with relatively short times to reduce below 5% of initial occupancy in the brain (<24 hours).

### Higher-functioning aptabots with affinity loss and affinity gain capacities

There are several advantages for a contrast agent to possess affinity gain capacities in addition to affinity loss capacities. **Figure 3D** illustrates a contrast agent with a single arm that can inducibly gain affinity then lose affinity for a target; the imaging advantage to this agent, when also equipped with necessary functionalization to bypass the BBB, is that it retains extra stealth from immune recognition until it is in the brain and the researcher chooses inducibly to release the protector strands. Another advantage is that if it targets a receptor that exists within and outside of the brain, then this agent can be restricted from having affinity for that receptor until after bypassing the BBB. More advantages still exist for an agent with multiple distinct arms similar to those in figure 3D; this agent can be programmed to bind new targets after releasing from prior ones. This benefit saves a researcher from both waiting and checking for systemic clearance, as well as from needing to construct and add new contrast agents.

In order for a contrast agent only to gain aptamer-facilitated target affinity upon molecular induction, the strand that folds into the facilitating aptamer must initiate the process in a shielded state, hybridized to complementary DNA. Figure 4 illustrates two options for how this shielding may be facilitated, with either a single protector strand or multiple protector strands. In both cases, the protector strands contain toeholds, and are removed after substitute strands induce displacement reactions. The downside to using a single protector strand is that the substitute strand contains the full aptamer sequence, and may therefore fold before reaching its target. Using multiple protector strands, each with a toehold sequence, circumvents this issue: each substitute strand complementary to a protector strand contains only a fraction of the strand sequence of the strand that will fold into an aptamer, without folding itself into an aptamer in solution.

**Figure 4.**
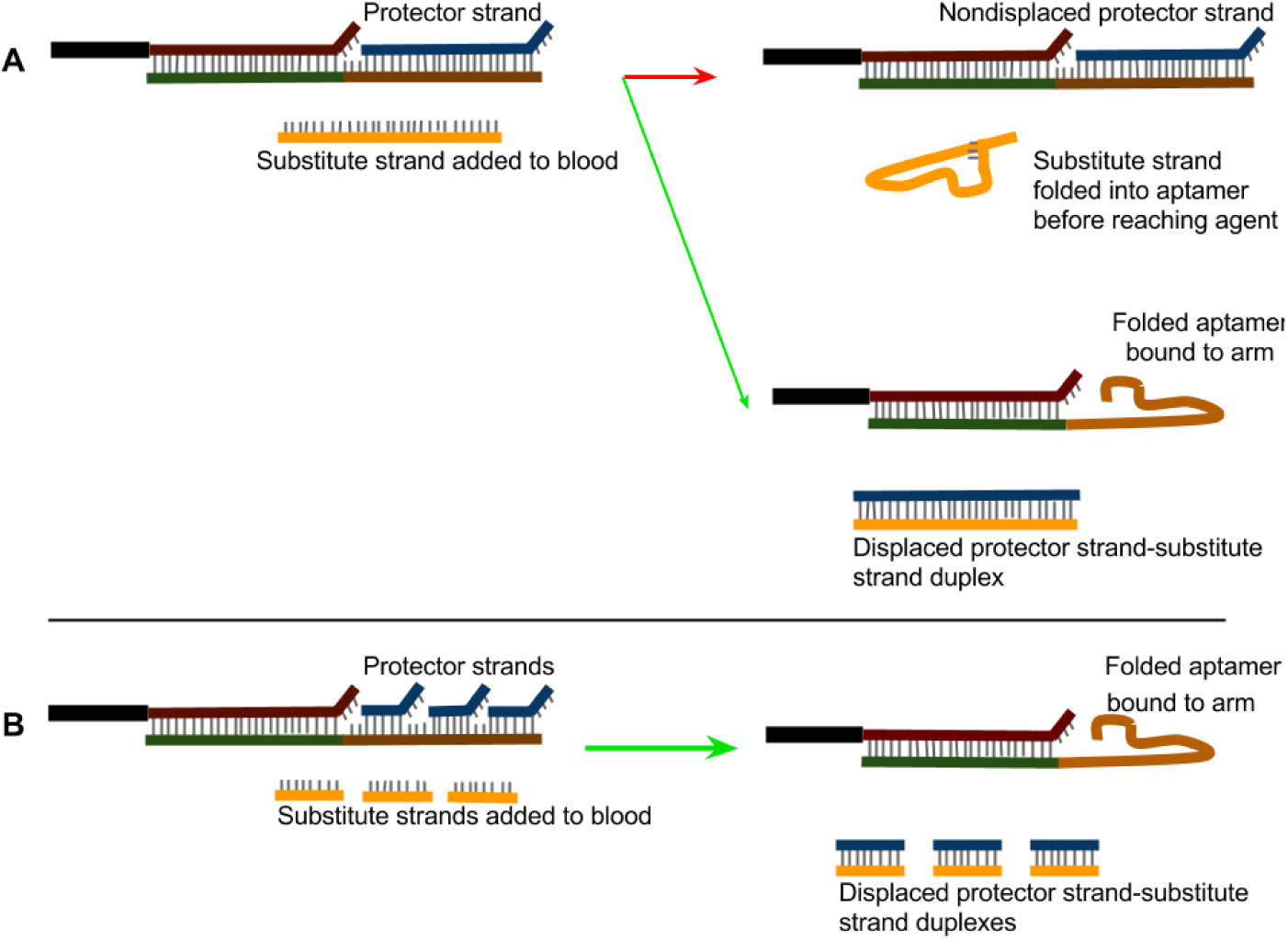
Comparison between outcomes when **(A)** shielding an aptamer-folding strand with a single protector strand and when **(B)** shielding an aptamer-folding strand with multiple protector strand.

A fully functional affinity-switching contrast agent is illustrated in Figure 5, with 5 distinct arm types: 4 for distinct imaging targeting, and 1 for RMT-targeting. In this illustration, where several stages of contrast agent function are shown, affinity loss for each prior imaging target is induced before affinity gain for each subsequent imaging target; however, these steps may be reversed or simultaneously conducted.

**Figure 5.**
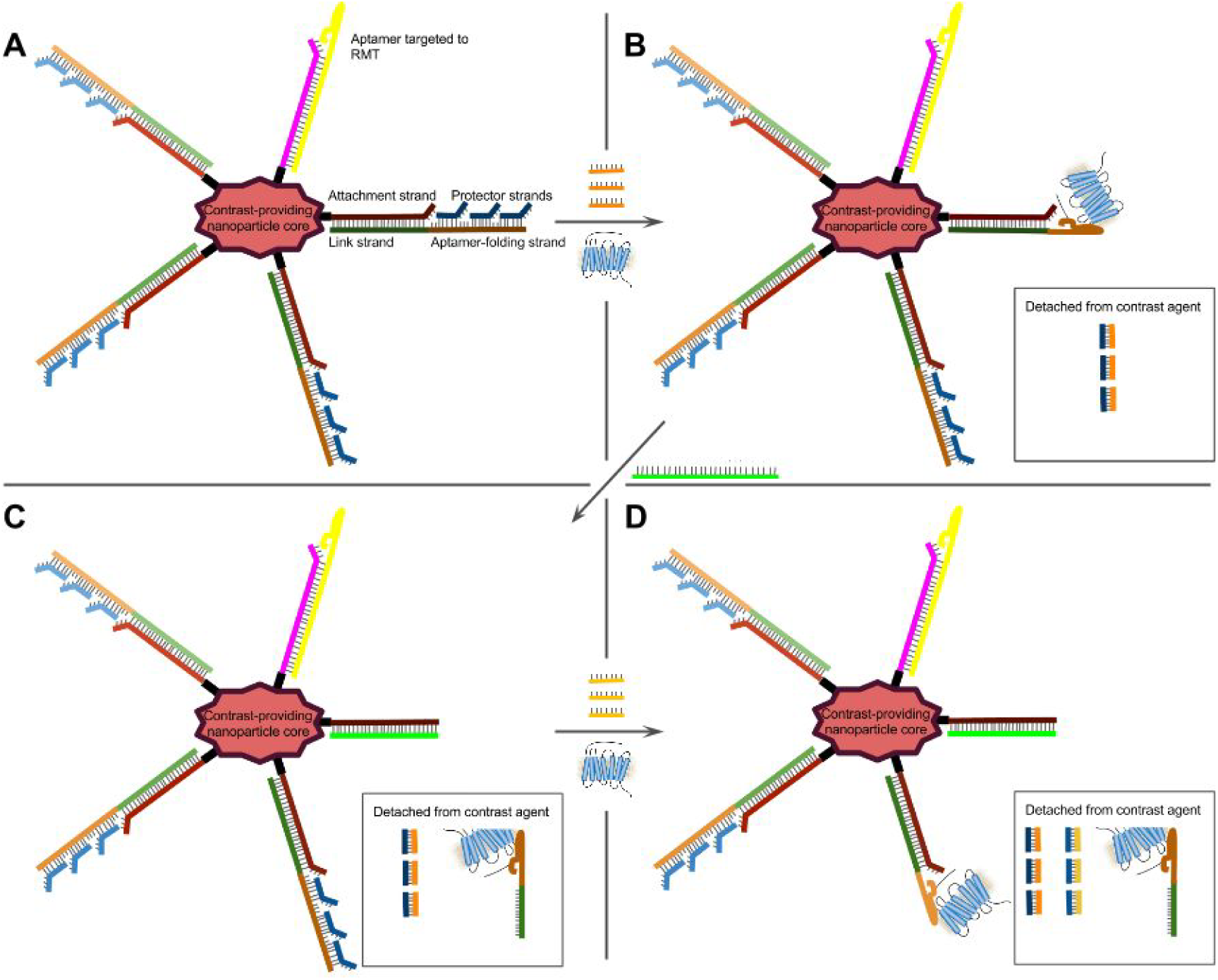
Affinity-switching contrast agent schematic. **A.** Contrast agent prior to any strand displacement reactions. Affinity strands (**orange**, in sequential fade) are each shielded by multiple bound toehold-containing protector strands (**blue**, in sequential fade) and are concatenated to link strands (**green**, in sequential fade) that are bound to toehold-containing attachment strands (**red**, in sequential fade). Attachment strands are conjugated to a contrast-providing nanoparticle (**pink**). An additional arm with an attachment strand (**magenta**) hybridized to a strand whose terminus is pre-emptively folded into an RMT-targeting aptamer (**yellow**) encourages BBB transport of the agent. The toehold on the magenta attachment strand allows inducible removal of the RMT-targeting aptamer, which may or may not have functional benefit. **B.** Contrast agent after the first set of strand displacement reactions, bound to the first transmembrane protein target. **C.** Contrast agent after the first two sets of strand displacement reactions. **D.** Contrast agent after the first three sets of strand displacement reactions, bound to the second transmembrane protein target.

If affinity-switching contrast agents, as illustrated in Figure 5, meet the following conditions, they will function as desired: a) reach their targets, b) are adequately reached by substitute strands, and c) have substantially long toeholds for rapid displacement kinetics and strong displacement-favoring equilibria. The illustrated agent will sequentially provide contrast for four different imaging targets with simple induction procedures. Agents can be manufactured to provide sequential contrast for up to ten different imaging targets or more, given the right synthesis conditions (**Table 3**). The use of affinity-switching agents provides a some temporal advantage over using releasable contrast agents with only single affinities, since 17-hour clearance times are still necessary to remove at least 95% of released agents from the brain, and such clearance is unnecessary if an agent alternatively acquires affinity for a new imaging target within the brain. Approximately 6.25 hours are necessary between imaging procedures when switching an agent from one affinity to the next using strand displacement, whereas there is a minimum 23.25 hour delay when clearing one agent from the system and adding another between imaging procedures. The net consequences of these differences, when sequentially imaging multiple targets in a single organism, can be visualized in Figure 6. If one considers that non-releasable agents have a minimum clearance delay of 68 hours (with low affinity for their targets) and a maximum clearance delay of years (with higher affinity for their targets), the imaging times illustrated in this figure all show a clear advantage over what would be required, in many cases impossible, with non-releasable or non-multi-affinity agents.

**Table 2.** Projected total time (hours) to image target populations of a given number of target types, with sequential populations of contrast agents that each have a given number of potential affinities. All contrast agents are considered to be inducibly releaseable from targets. 3 hours are given for contrast agent diffusion across the BBB. 1 hour is given for imaging. 3 hours are given for substitute strand diffusion across the BBB together with 0.25 hours for 95% reaction completion (given 6 nucleotide toehold lengths). 17.29 hours are given for 95% agent clearance after all affinities of a given agent population have been used: this would be accurate given 4hour brain halflives of the released agents, although the brain halflives of released agents may be longer. Longer values increase imaging time ratios between when using contrast agents with fewer potential affinities and when using contrast agents with more potential affinities.

|  | 1 potential affinity per contrast agent | 4 potential affinities per contrast agent | 10 potential affinities per contrast agent |
| --- | --- | --- | --- |
| 1 imaging target | 4 | 4 | 4 |
| 4 imaging targets | 77.61 | 16.75 | 16.75 |
| 10 imaging targets | 224.84 | 82.83 | 42.25 |
| 20 imaging targets | 470.22 | 165.9 | 105.04 |

**Table 3.** Predicted percentages of 13 nm gold nanoparticles conjugated to thiolated attachment strands of a full typal diversity, following conjugation with equimolar excess concentrations of each thiolated attachment strand type in solution, depending upon different buffer conditions corresponding to different adsorption levels of thiolated attachment strands. For nanoparticles of radius r with a similar shape to those used by Zhang et al, DNA adsorption levels may be predicted to equal those in column 2 times the quantity r^2^/13^2^, and other values are shifted accordingly.

| [NaCl] (mM) during conjugation of thiolated attachment strands to citrate-capped AuNPs; pH during conjugation | Approximate number of thiolated attachment strands adsorbed per AuNP (Zhang X 2012) | Percent fully diversified AuNPs given 2 thiolated attachment strand types in equimolar excess concentrations in conjugation solution | Percent fully diversified AuNPs given 5 thiolated attachment strand types in equimolar excess concentrations in conjugation solution | Percent fully diversified AuNPs given 11 thiolated attachment strand types in equimolar excess concentrations in conjugation solution |
| --- | --- | --- | --- | --- |
| 30; 7 | 30 | **100** | **99** | **49** |
| 30; 3 | 40 | **100** | **100** | **77** |
| 100; 7 | 50 | **100** | **100** | **91** |
| 100; 3 | 65 | **100** | **100** | **98** |
| 300; 7 | 55 | **100** | **100** | **94** |
| 300; 3 | 75 | **100** | **100** | **99** |

**Figure 6.**
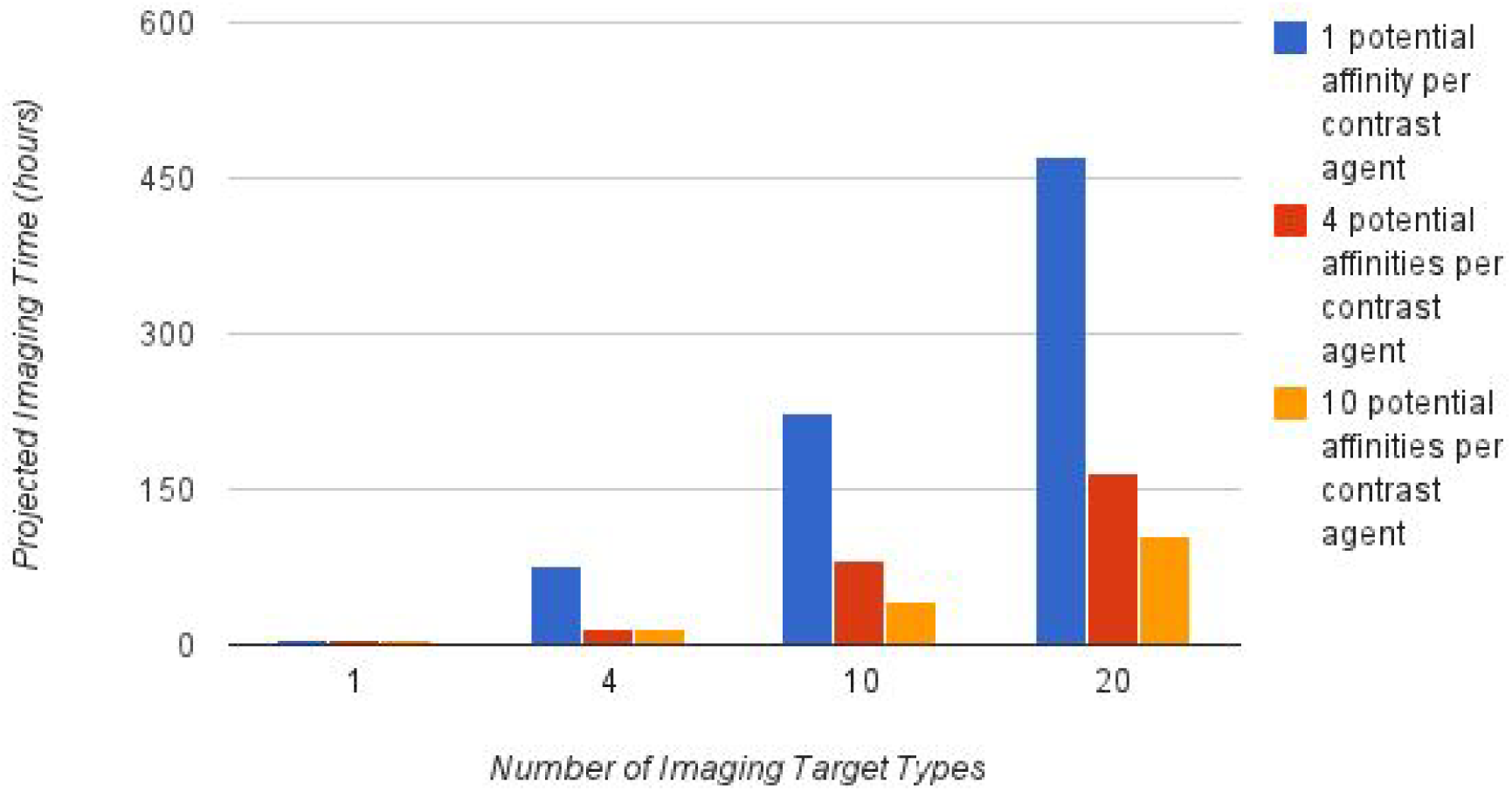
Bar graph of total time (hours) to image target populations of a given number of target types, with sequential populations of contrast agents that each have a given number of potential affinities. All contrast agents are considered to be inducibly releasable from targets. Bar graph based on values in Table 2.

### Construction Considerations for Multi-Stage Aptabots

Construction of multi-stage/multi-affinity aptabots requires important DNA adsorption considerations, in order to ensure a fully diverse quota of arms conjugates to each of a large portion of nanoparticles. The portion of nanoparticles to which a fully diverse quota of arms conjugates, given random conjugating conditions, is dependent on the total number of arms conjugated to each nanoparticle. Assuming there are m distinct arm types, and that n arms conjugate to a nanoparticle (n>m), each of a random type, random conjugating conditions can be enforced by using equimolar concentrations of each arm type in solution during the conjugation step, and by using a conjugation method that does not favor a particular arm type and that gives rise neither to cooperative nor to noncooperative binding properties between arms of particular type pairs.

The probability that at least one arm of each type binds to the particle is equal to 1 minus the probability that no arm of at least one type binds to the particle. The inclusion-exclusion principle yields the following equation for the probability that no arm of at least one type binds to the particle:

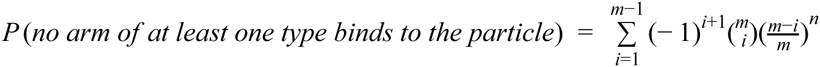

In the summand, the product 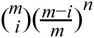 is equal to the number of unique combinations of *i* arm type choices times the probability that no arm of any type amongst a given combination of *i* arm type choices binds to the particle. Using the inclusion-exclusion principle to alternate adding and subtracting these products, we can account for any unique possibility of failures to bind arms from a selection of given arms exactly once. The probability that at least one arm of each type binds to the particle is equal to 1. This sum, and the percentage of particles in a population that binds at least one arm of each type is equal to the probability that at least one arm of each type binds to a single particle.

When conjugating thiolated DNA to citrate-capped 13 nm gold nanoparticles (AuNPs), DNA adsorption levels have been measured in various buffer conditions (Zhang X 2012). With higher pH and lower sodium chloride concentrations, an average of approximately 20 DNA molecules were adsorbed per nanoparticle; with lower pH and higher sodium chloride concentrations, an average of approximately 75 DNA molecules were adsorbed per nanoparticle. Higher adsorption levels are important when attempting to conjugate at least one of each arm type to a large percentage of nanoparticles, when a large variety of arm types need to be attached. Similar to the conditions used in the diversity-determining step of nanoparticle conjugation and functionalization, attachment strands can be thiolated and then conjugated to nanoparticles. According to hybridization protocols, the remainder of each DNA arm can subsequently be hybridized to the specific attachment strand for which it has affinity (Kim 2010). Table 3 displays the percentage of particles that absorb at least 1 of each attachment strand species given a certain number of thiolated attachment strand species in equimolar concentrations in the conjugation solution when using different conjugation conditions of thiolated attachment strands to nanoparticles that give rise to various adsorption levels per nanoparticle.

When conjugating more distinct varieties of attachment strands to nanoparticles, choosing conjugation conditions that maximize adsorption levels grows increasingly important, to ensure a high proportion of contrast agents have at least one of attachment strand species. DNA adsorption levels may vary when one uses differently sized and shaped nanoparticles. Table 3 serves to indicate that for at least one nanoparticle species within the small size range, a conjugation approach and condition exists to reach adsorption levels that will guarantee a sufficiently high percentage of fully-functionalized contrast agents. It also indicates, however, that said conjugation approach and condition must be carefully selected in order to guarantee this percentage.

### Alternative uses: siRNA

There are numerous uses for variations of the technology. It can be used for mapping multiple submicron details of organs or organ systems in a living body, at fine resolutions and without the need to wait for long agent clearance times. The brain is the most challenging such organ, as imaging submicron details of the *in vivo* brain requires BBB transversal of agents and provision of contrast for skull-penetrating high-resolution methods. But reaching imaging goals for any organ or organ system may be greatly accelerated by technology that enables inducible affinity-loss and affinity-gain imaging agents.

One less obvious use for this technology is the inducible delivery of RNA interference to local cellular populations. Consider the agent illustrated in Figure 7. This agent migrates towards and binds specifically to cancer cell populations, and can be imaged to verify its localization to a known tumor in a system. Once its location is verified, RNA substitute strands can be added to the blood to release from the agent’s double-stranded RNA, which subsequently will be transported through cellular membranes and recognized by DICER, for the production of microRNA, provided the proper sequence tags are included. If this microRNA is targeted to inhibit important cell survival processes or inhibitors of apoptosis, the either necrosis or apoptosis can be locally induced. Proximal blood vessels providing the tumor with nutrients may also be targeted. The agent itself then can be inducibly released from the vicinity to expedite its clearance. This approach can be generalized towards the delivery of any tissue behavior-modifying RNAi to any region of tissue expressing unique surface markers for a variety of other uses.

**Figure 7.**
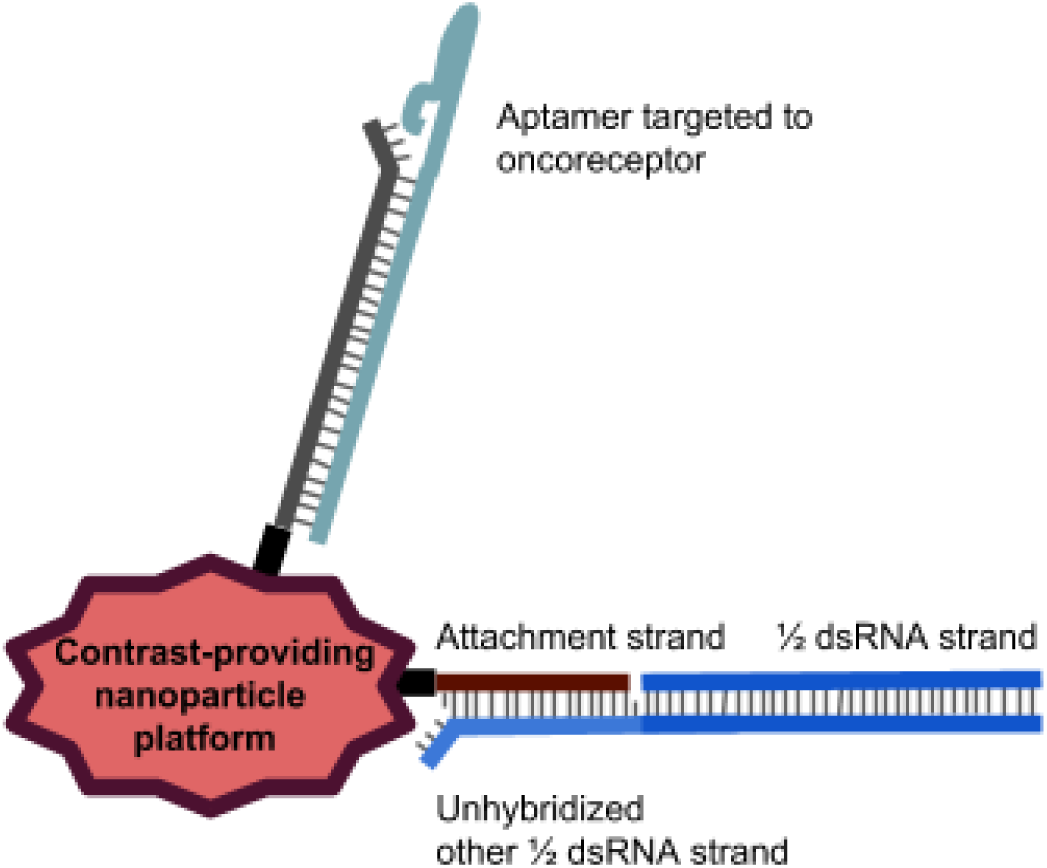
RNAi-to-tumor delivery agent.

## Discussion

The contrast agents proposed in this paper can be constructed with ease. If gold nanoparticles (AuNPs) be used as the nanoparticle cores, only a few modifications to the protocol developed and published Kim et al (2010) need to be applied to construct the contrast agents illustrated here. These alterations would involve initially conjugating equimolar concentrations of distinct thiolated attachment strands, including toehold sequences at the 3’ ends of attachment strands, and hybridizing protector strands to unfolded aptamer-folding strands at temperatures decreasing from denaturing to annealing ranges prior to hybridizing link strands. Should superparamagnetic iron oxide nanoparticles (SPIONs) be used as the nanoparticle cores, a few adjustments to the protocol developed and published in 2008 by Wang et al. for such conjugation are necessary. These changes are similar to those listed above, with the additional adjustment of conjugating aminated attachment strands to carboxylated SPIONs instead of aminated aptamer-folding strands.

The agents proposed in this paper are designed to provide contrast for specific brain targets *in vivo*. Several agents similar to those outlined in this paper have been constructed and used to provide contrast for specific brain targets or targets in other organs *in vivo*. Citrate-capped gold nanoparticles conjugated to thiolated ssDNA strands, hybridized to aptamer-concatenated link strands, have been utilized for targeted CT imaging of and drug delivery to prostate cancer cells (Wang 2008). Superparamagnetic iron oxide nanoparticles conjugated via peptide bonds to aptamers have been utilized for targeted MRI imaging of and drug delivery to prostate cancer cells (Kim 2010). Nanoprobes dual-targeted to bypass the blood-brain-barrier with RMT-targeting antibodies have been used for targeted MRI imaging of specific glioblastoma proteins (Ni 2014).

In regards to constructing the specific contrast agents illustrated in this paper, several investigators have identified specific aptamers that could prove useful. Recent promising publications describe an RMT-targeting aptamer (Cheng 2003) as well as an aptamer targeted to the endothelial regulatory protein PIGPEN, which is overexpressed in brain tumor microvessels (Blank 2001). With concern to synaptic targets, Ferapontova et al. developed and utilized aptamers targeted to dopamine (Cheng 2013, Blank 2001, Farjami 2012).

The only salient challenges to creating and fully applying contrast agents as designed and illustrated in this paper are A) identifying aptamers with adequate specificity and affinity for brain proteins of imaging interest and B) ensuring appropriate reagents cross the BBB. (A) simply seems unaddressed in literature. Antibodies and chemical ligands have provided adequate affinity for imaging targets of interest with contrast agents used in imaging following slicing and staining. Now that higher resolution non-destructive imaging is a greater possibility with the emergence of finer imaging procedures, there are strong motivations for using aptamers to provide affinity for imaging targets, such as a lower cost for broader-scale imaging and inducible affinity gain/loss via strand exchange. (B) may be adequately addressed on multiple reagents—contrast agents and substitute strands for inducing strand exchange—by using the RMT-targeting aptamer (Cheng 2013).

As suggested, the benefits to adequately constructing and utilizing the proposed contrast agents are tremendous. First, non-destructive imaging of the rodent or human brain can enable better understanding of not only immediate brain architecture associated with given healthy or diseased states, but also the change over time of brain architecture associated with the development and/or progression of these states in a single organism’s brain, which is something destructive imaging could never accomodate. There are myriad diseases towards which this technology could be applied. Flynn et al discovered shifts in myelination and oligodendrocyte-associated protein levels associated with the development of schizophrenia in postmortem rodents (Flynn 2003). With the contrast agents we propose, levels and distributions of these oligodendrocyte-associated proteins could be monitored over time in single living organisms. These organisms could possibly even include humans if safe contrast-providing particle cores such as feraheme (ferumoxytol) are used (Lu 2010, Castaneda 2011).

Abnormalities in cerebral brain structure have historically been observed in juvenile myoclonic epilepsy, but specific circuit arrangement abnormalities associated with these structural abnormalities—and the correlated development of circuit arrangement abnormalities associated with that of these structural abnormalities—have yet to be deciphered due to the absence of safe, specifically targeted brain imaging agents (Woermann 1999, O’Muircheartaigh 2011). Many other neurological diseases exist for which high-resolution time course imaging of brain architecture would an be invaluable tool to understand their pathology. According to the foci of current literature, schizophrenia and epilepsy are potential choices, as the descriptions of neural architectural changes must be complete in order to prescribe intervention or treatment. Furthermore, observing changes in a single healthy brain’s architecture over time could be invaluable towards understanding natural brain development, particularly changes in circuit structure associated with major phases of learning and maturation.

Second, the affinity-switching capacities of the contrast agents in this paper, together with their capacity to non-destructively provide contrast for high-res imaging methods, allow the imaging of multiple distinct circuitry facets in a healthy or diseased brain in acceptable timeframes for map overlays to be of value. Thus far, no contrast agents for imaging *in vivo* the architecture of any organ, much less a brain, have been able to promise every nuance of this capacity - high enough resolution to see circuit details, as well as quick subsequent imaging of multiple distinct facets of circuitry. Prior agents have been developed that have inducible affinity or contrast-providing activation and/or inactivation *in vivo*, but affinity or contrast switching of these other agents has relied on either pH-induced shifts of structural conformation or oxidation events that change agent structure. The ability of an agent to inducibly shift affinity or contrast in response to either pH or oxidizing agents is invaluable: it ensures an agent is only active in tissue of interest (e.g. cancer tissue) (Lowe 2004); on the other hand, it cannot be used functionally to manually to induce multiple distinct shifts in affinity or to induce any shifts in affinity without changing growth conditions of imaged tissue. Regarding expedited contrast agent clearance, biodegradable contrast agents do exist but they are slow to degrade and/or provide contrast for lower-resolution imaging techniques (Tam 2010). Contrast agents for PET degrade more quickly, and thus can be used for imaging multiple distinct architectural facets of a single organ in short timeframes (Reiner 2012), but the resolution for PET is also lower than that which can be acquired using MRI or X-ray CT (Moses 2011).

Imaging multiple architectural details of a single organism’s brain without harming the organism can enable better understanding of fully detailed circuit structure and changes associated with disease progression, neurological development, or specific learning events. This understanding, garnered for both a healthy and diseased organism, would both be valuable and far beyond what present science has been able to obtain. Ultimately, these contrast agents for imaging an arbitrary number of distinct architectural aspects within a single living brain at high resolutions can help solve a major challenge in neurobiology, connectome mapping. Recently, a map of the rodent connectome at a resolution of 1 micron was published, but the current state of technology required fine tissue slicing and staining (Oh 2014). *In vivo* connectome mapping could be conducted at similar resolutions with proper BBB-transversing, inducibly affinity-switching contrast agents. In general, functional agents such as those in this paper would make all imaging and diagnostics more efficient, inexpensive, and effective.

### Selected Materials and Methods Protocols for Aptabot Construction and Testing

#### 1. Identification of aptamers specific protein targets of interest

Purified protein is sourced commercially or overexpressed in cell lines and purified using current published microbiological techniques. In general it’s best to choose a protein that has been developed in as close to the in vivo model one hopes to image in. Thus, human cell line expression is prefered, and the use of whole cells, or lipid rafts for membrane bound proteins should give the best result when raising aptamers.

Purified proteins, or cell lines with overexpressed membrane bound proteins can then be used to select for or against aptamers to specified targets during a SELEX development process. The SELEX process can also be outsourced or performed via current published protocols. The final product is optimized aptamer sequences and folding buffer conditions for your target of interest.

Biotinylated versions of optimized aptamer sequences are ordered from oligomer producers such as Trilink Biotech. Biotinylated oligomers are used to verify kinetics, affinity levels, and specificity of each folded aptamer to its target via surface plasmon resonance (SPR) or other kinetic assay.

#### 2. Attachment strand functionalization of gold nanoparticles

For each desired nanoparticle arm, attachment strands may be ordered with 5’ thiol groups and with 3’ toehold stretches (either GGGGGG or CCCCCC). Citrate-capped nanoparticles and thiolated attachment strands in equimolar excess concentrations are conjugated in the 300 mM NaCl, pH 3.0. (Zhang X 2012).

#### 3. Attachment strand functionalization of iron oxide nanoparticles

For each desired nanoparticle arm, attachment strands are ordered with 5’ thiol groups and with 3’ toehold stretches (either GGGGGG or CCCCCC). Superparamagnetic iron oxide nanoparticles (SPIONs) may be constructed, carboxylated, and conjugated to 5’ aminated attachment strands in equimolar excess using EDC/NHS according to the protocols developed by Wang et al. (Wang 2008).

#### 4. Construction of aptamer arms and of substitute strands

Synthesis of subsitute strands or uninduced affinity arms is straightforward. For uninduced affinity-providing arms, such as the yellow arm in **Figure 5** providing RMT targeting, one simply synthesizes the affinity (aptamer-folding) strand sequence concatenated link strand sequence. Substitute strands are ordered as full complements to their target protector ore attachment strands.

For induced affinity-providing arms, synthesis involves ordering oligonucleotides with the affinity strand sequence concatenated at the 3’ end to the link strand sequence. Carefully chosen protector strands complementary to disjoint stretches of the affinity strand sequence concatenated to 3’ toehold regions must also be synthesized. To select protector strand oligonucleotides one must identify sequences of maximum length in the affinity strand that, when ordered separately as substitute strands each concatenated to 5’ toeholds (either GGGGGG or CCCCCC), do not have the tendency to fold into undesirable secondary structures such as hairpins or loops.

Protector strands and affinity strands are annealed by mixing protector strands and affinity strand-link strand concatamers in equimolar concentrations in solution and then elevating temperatures to denaturing ranges to induce aptamers to unfold. Fully complementary binding is the first to have a negative Gibbs free energy change associated with each successively higher concentration ratio of duplexes to free strands as temperature decreases. As such, slowly lowering the temperature towards annealing ranges favors the formation exclusively of duplexes with the greatest level of complementarity. This ensures that protector strands anneal to affinity strands and that affinity strands do not refold into aptamers.

#### 5. Hybridization of aptamer arms to attachment strand-functionalized nanoparticles

Completed arms are hybridized to attachment strand-functionalized nanoparticles when hybridizing aptamer strand-link strand concatenations to ONT-functionalized nanoparticles to functionalize these nanoparticles for prostate cancer cell targeting (Kim 2010).

#### 6. Testing functional aptamer hybridization and inducible affinity loss/gain capacities

Adequacy of conjugation methods and stability of attachment strand-mediated conjugations of nanoparticles to link-strand-aptamer concatenations can be firstly measured by testing the construction and stability of simplest-case conjugates, where nanoparticles are functionalized with single attachment strand species and subsequently hybridized to single arm species without protector strands (i.e., with pre-emptively exposed and folded aptamers). Testing the construction, affinity, and specificity of these nanoparticle-aptamer conjugates can be initiated by purifying conjugates, and then by mixing combinatorially in buffer solution that emulates BECF buffer conditions: i) complete conjugates, free nanoparticles, or attachment strand-functionalized nanoparticle controls without aptamers, with ii) purified target proteins or purified non-target proteins. Such tests can proceed with the re-purification of nanoparticles and the conduction of ELISA binding assays for the presence of target proteins on nanoparticles. Testing the stability of these conjugates can be enacted by subjecting conjugates to varying conditions, and then by repeating the tests for construction and affinity.

Next, tests for inducible affinity loss capacity can be conducted via two methods. The first is treating conjugates with substitute strands for link strands, and subsequently testing affinity and specificity (as described above) for treated vs. untreated conjugates. The second is mixing conjugates with purified target proteins before treating one population with substitute strands for link strands and leaving another untreated, then re-purifying conjugates and afterwards conducting ELISA binding assays for the presence of target proteins on treated vs. untreated conjugates.

Tests for inducible gain in affinity can be conducted by constructing conjugates, as in Figure 4, and then treating conjugates as described above for treated vs. untreated conjugates.

Finally, testing for adequate construction and stability of more complex nanoparticles, as well as for sequential affinity gain and loss capacities of these nanoparticles, can be arranged by making more complex nanoparticles according to the above protocols. One population of particles can be constructed with preemptively exposed aptamer arms, and another with shielded arms. From here, variations of the tests described above can also be conducted.

The above tests involve the verification of particle construction, affinity, and inducible affinity/loss capacities in buffer conditions similar to those in BECF, and as well as the verification of particle stability in buffer conditions, similar to those in blood plasma. Following these tests, *in vitro* functional tests can be conducted instead of using ELISA binding assays. By employing assays that compare the extent of co-localization for binding, we can examine the difference between a) the extent of co-localization of agents with fluorochrome-tagged target proteins and b) the extent of co-localization of agents with fluorochrome-tagged control proteins that are not targets of interest, to ensure co-localization does not occur with arbitrary protein clusters. Imaging in this case utilizes fluorescence to visualize target protein localization and then the imaging method for which the nanoparticle provides optimal contrast to visualize agent localization.

## Supplementary Mathematics

If a system is monocompartmental, and a free contrast agent B has a clearance-induced half-life in the system of n days,

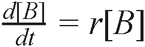

where t is time in days and r=1/n*ln(0.5).

Now let contrast agent B associate with targets A. Let K_D_ be the dissociation constant, k1 be the association reaction rate constant, k2 be the dissociation reaction rate constant, and [B_total_] be the total concentration [AB] + [B]. In this case,

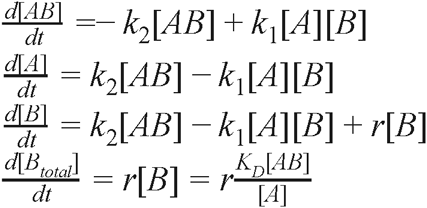

[A] is in excess, so K_D_/[A] may be treated as a constant, 1/L.

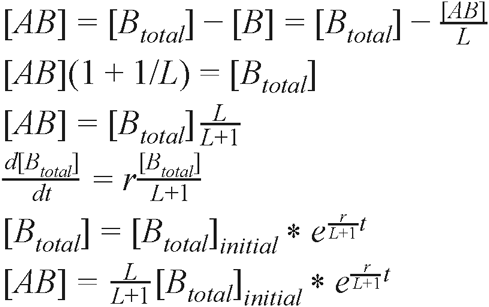

For half-life T_1/2_ of [AB],

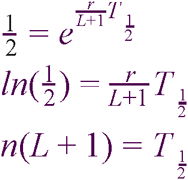

## References

Abbott, N Joan, Adjanie A K Patabendige, Diana E M Dolman, Siti R Yusof, and David J Begley. “Structure and Function of the Blood-Brain Barrier.” Neurobioloav of Disease 37, no. 1 (January 2010): 13–25.

Alivisatos, A. P., A. M. Andrews, et al. “Nanotools for Neuroscience and Brain Activity Mapping.” Acs Nano 7, no. 3 (2013): 1850–1866.

Arvantis, C.D., M.S. Livingstone, N. Vykhodtseva, and N. McDannold. “Controlled Ultrasound-Induced Blood-Brain Barrier Disruption Using Passive Acoustic Emissions Monitoring.” PLoS ONE 7 (2012) e45783.

Asharani, P. V., Y. Lianwu, et al. “Comparison of the toxicity of silver, gold and platinum nanoparticles in developing zebrafish embryos.” Nanotoxicoloav 5, no. 1 (2011): 43–54.

Beekman FJ, van der Have F, Vastenhouw B, van der Linden AJ, van Rijk PP, Burbach JP, Smidt MP. “U-SPECT-I: a novel system for submillimeter-resolution tomography with radiolabeled molecules in mice.” J Nucl Med 46, no. 7 (July 2005): 1194–200.

Blank, M, T Weinschenk, M Priemer, and H Schluesener. “Systematic Evolution of a DNA Aptamer Binding to Rat Brain Tumor Microvessels. Selective Targeting of Endothelial Regulatory Protein Pigpen.” The Journal of Biological Chemistry 276, no. 19 (May 11, 2001): 16464–68.

Burbulis, I., K. Yamaguchi, et al. “Quantifying small numbers of antibodies with a 'near-universal' protein-DNA chimera.” Nat Methods 4, no. 12 (2007): 1011–1013.

Cao, Z., R. Tong, et al. “Reversible cell-specific drug delivery with aptamer-functionalized liposomes.” Anoew Chem Int Ed Engl 48, no. 35 (2009): 6494–6498.

Cassette, Elsa, Marion Helle, Lina Bezdetnaya, Frédéric Marchal, Benoit Dubertret, and Thomas Pons. “Design of New Quantum Dot Materials for Deep Tissue Infrared Imaging.” Advanced Drug Delivery Reviews 65, no. 5 (May 2013): 719–31.

Castaneda, R. T., A. Khurana, et al. “Labeling stem cells with ferumoxytol, an FDA-approved iron oxide nanoparticle.” J Vis Exp 57 (2011): e3482.

Cavalieri, F., M. Zhou, et al. “Methods of preparation of multifunctional microbubbles and their in vitro / in vivo assessment of stability, functional and structural properties.” Curr Pharm Des 18, no. 15 (2012): 2135–2151.

Chang, J. C., O. Kovtun, et al. (2012). “Labeling of neuronal receptors and transporters with quantum dots.” Wiley Interdisciplinary Reviews: Nanomedicine and Nanobiotechnology 4(6): 605–619.

Chen, Y. C., W. R. Galpern, et al. “Detection of dopaminergic neurotransmitter activity using pharmacologic MRI: correlation with PET, microdialysis, and behavioral data.” Maon Reson Med 38, no. 3 (1997): 389–398.

Cheng, Congsheng, Yong Hong Chen, Kim A Lennox, Mark A Behlke, and Beverly L Davidson. “In Vivo SELEX for Identification of Brain-Penetrating Aptamers.” Molecular Therapy. Nucleic Adds 2 (2013): e67.

Cho, Hoonsung, David Alcantara, Hushan Yuan, Rahul A. Sheth, Howard H. Chen, Peng Huang, Sean B. Andersson, David E. Sosnovik, Umar Mahmood, and Lee Josephson. “Fluorochrome-Functionalized Nanoparticles for Imaging DNA in Biological Systems.” ACS Nano 7, no. 3 (March 26, 2013): 203–211.

Chung, K., J. Wallace, et al. “Structural and molecular interrogation of intact biological systems.” Nature 497, no. 7449 (2013): 332–337.

De Rosa, G. and K. I. La Rotonda. “Nano and microtechnologies for the delivery of oligonucleotides with gene silencing properties.” Molecules 14, no. 8 (2009): 2801–2823.

Dreaden, Erik C, Lauren A Austin, Megan A Mackey, and Mostafa A El-Sayed. “Size Matters: Gold Nanoparticles in Targeted Cancer Drug Delivery.” Therapeutic Delivery 3, no. 4 (April 2012): 457–78.

Farde, Lars, Håkan Hall, Stefan Pauli, and Christer Halldin. “Variability in D2-Dopamine Receptor Density and Affinity: A PET Study with [11C]raclopride in Man.” Synapse 20, no. 3 (July 1, 1995): 200–208.

Farjami, Elaheh, Rui Campos, Jesper S. Nielsen, Kurt V. Gothelf, Jørgen Kjems, and Elena E. Ferapontova. “RNA Aptamer-Based Electrochemical Biosensor for Selective and Label-Free Analysis of Dopamine.” Analytical Chemistry 85, no. 1 (January 2, 2013): 121–28.

Ferreira, C. S., M. C. Cheung, et al. “Phototoxic aptamers selectively enter and kill epithelial cancer cells.” Nucleic Acids Res 37, no. 3 (2009): 866–876.

Ferreira, C. S., K. Papamichael, et al. “DNA aptamers against the MUC1 tumour marker: design of aptamer-antibody sandwich ELISA for the early diagnosis of epithelial tumours.” Anal Bioanal Chem 390, no. 4 (2008): 1039–1050.

Flynn, S. W., D. J. Lang, et al. “Abnormalities of myelination in schizophrenia detected in vivo with MRI, and post-mortem with analysis of oligodendrocyte proteins.” Mol Psychiatry 8, no. 9 (2003): 811–820.

Gardner, D., H. Akil, et al. “The Neuroscience Information Framework: A Data and Knowledge Environment for Neuroscience.” Neuroinformatics 6, no. 3 (2008): 149–160.

Geraldes, C. F. and S. Laurent. “Classification and basic properties of contrast agents for magnetic resonance imaging.” Contrast Media Mol Imaging 4, no. 1 (2009): 1–23.

Geszke-Moritz, Malgorzata, and Michal Moritz. “Quantum Dots as Versatile Probes in Medical Sciences: Synthesis, Modification and Properties.” Materials Science & Engineering. C. Materials for Biological Applications 33, no. 3 (April 1, 2013): 1008–21.

Gulati, M., D. S. Chopra, et al. “Patents on brain permeable nanoparticles.” Recent Pat CNS Drug Discov 8, no. 3 (2013): 220–234.

Hayworth, K., N. Kasthuri, et al. “Automating the Collection of Ultrathin Serial Sections for Large Volume TEM Reconstructions.” Microscopy and Microanalysis 12, Supplement S02 (2006): 86–87.

Hellebust, A. and R. Richards-Kortum. “Advances in molecular imaging: targeted optical contrast agents for cancer diagnostics.” Nanomedicine 7, no. 3 (2012): 429–445.

Helmstaedter, K., K. L. Briggman, et al. “Connectomic reconstruction of the inner plexiform layer in the mouse retina.” Nature 500, no. 7461 (2013): 168–174.

Javasena, S. D.. “Aptamers: an emerging class of molecules that rival antibodies in diagnostics.” Clin Chem 45, no. 9 (1999): 1628–1650.

Jiao, P F, H Y Zhou, L X Chen, and B Yan. “Cancer-Targeting Multifunctionalized Gold Nanoparticles in Imaging and Therapy.” Current Medicinal Chemistry 18, no. 14 (2011): 2086–2102.

Jing, Meng, and Michael T. Bowser. “A Review of Methods for Measuring Aptamer-ProteinEquilibria.” Analytica Chimica Acta 686, no. 1-2 (February 7, 2011): 9–18.

Kim, Dongkyu, Yong Yeon Jeong, and Sangyong Jon. “A Drug-Loaded Aptamer-Gold Nanoparticle Bioconjugate for Combined CT Imaging and Therapy of Prostate Cancer.” ACS Nano 4, no. 7 (July 27, 2010): 3689–96.

Kim, Dongkyu, Sangjin Park, Jae Hyuk Lee, Yong Yeon Jeong, and Sangyong Jon. “Antibiofouling Polymer-Coated Gold Nanoparticles as a Contrast Agent for in Vivo X-Ray Computed Tomography Imaging.” Journal of the American Chemical Society 129, no. 24 (June 1, 2007): 7661–65.

Kopec, C. D., E. Real, et al. “GluR1 links structural and functional plasticity at excitatory synapses.” J Neurosci 27, no. 50 (2007): 13706–13718.

Le Novere, N.. “MELTING, computing the melting temperature of nucleic acid duplex.” Bioinformatics 17, no. 12 (2001): 1226–1227.

Lee, J. H.. “Conjugation approaches for construction of aptamer-modified nanoparticles for application in imaging.” Curr Top Med Chem 13, no. 4 (2013): 504–512.

Lee, Nohyun, Seung Hong Choi, and Taeghwan Hyeon. “Nano-Sized CT Contrast Agents.” Advanced Materials (Deerfield Beach, Fla.) 25, no. 19 (May 21, 2013): 2641–60.

Li, S., J. Jiang, et al. “PEGylation of protein-based MRI contrast agents improves relaxivities and biocompatibilities.” J Inoro Biochem 107, no. 1 (2012): 111–118.

Liu, H., H. Wang, Y. Xu, M. Shen, J. Zhao, G. Zhang, and X. Shi. “Synthesis of PEGylated low generation dendrimer-entrapped gold nanoparticles for CT imaging applications.” Nanoscale 6 (2014): 4521–4526.

Lowe, Mark P. “Activated MR Contrast Agents.” Current Pharmaceutical Biotechnology 5, no. 6 (December 1, 2004): 519–28.

Lu, M., et al. “FDA review of ferumoxytol (Feraheme) for the treatment of iron deficiency anemia in adults with chronic kidney disease.” Am J Hematol 85 (2010): 315–319.

Mamedov, I., S. Canals, et al. “In Vivo Characterization of a Smart MRI Agent That Displays an Inverse Response to Calcium Concentration.” ACS Chemical Neuroscience 1, no. 12 (2010): 819–828.

Mandeville, J. B., C. Y. Sander, et al. “A receptor-based model for dopamine-induced fMRI signal.” Neuroimaae 75 (2013): 46–57.

McCauley, T. G., N. Hamaguchi, et al. “Aptamer-based biosensor arrays for detection and quantification of biological macromolecules.” Anal Biochem 319, no. 2 (2003): 244–250.

Miller, J. M., J. E. Hutchison, et al. (2010). Functionalized nanoparticles and methods of forming and using same, Google Patents.

Montenegro, Jose-Maria, Valeria Grazu, Alyona Sukhanova, Seema Agarwal, Jesus M de la Fuente, Igor Nabiev, Andreas Greiner, and Wolfgang J Parak. “Controlled Antibody/(bio-) Conjugation of Inorganic Nanoparticles for Targeted Delivery.” Advanced Drug Delivery Reviews 65, no. 5 (May 2013): 677–88.

Morcos, S K. “Nephrogenic Systemic Fibrosis Following the Administration of Extracellular Gadolinium Based Contrast Agents: Is the Stability of the Contrast Agent Molecule an Important Factor in the Pathogenesis of This Condition?” The British Journal of Radiology 80, no. 950 (February 1, 2007): 73–76.

Moses, William W. “Fundamental Limits of Spatial Resolution in PET.” Nuclear Instruments & Methods in Physics Research. Section A. Accelerators. Spectrometers. Detectors and Associated Eouipment 648, Supplement 1 (August 21, 2011): S236–S240.

Nam, Jutaek, Nayoun Won, Jiwon Bang, Ho Jin, Joonhyuck Park, Sungwook Jung, Sanghwa Jung, Youngrong Park, and Sungjee Kim. “Surface Engineering of Inorganic Nanoparticles for Imaging and Therapy.” Advanced Drug Delivery Reviews 65, no. 5 (May 2013): 622–48.

Ni D, Zhang J, Bu W, Xing H, Han F, Xiao Q, Yao Z, Chen F, He Q, Liu J, Zhang S, Fan W, Zhou L, Peng W, and Shi J. “Dual-Targeting Upconversion Nanoprobes across the Blood Brain Barrier for Magnetic Resonance/Fluorescence Imaging of Intracranial Glioblastoma.” Acs Nano 8, no. 2 (2014): 1231–1242.

Niidome, T., M. Yamagata, et al. “PEG-modified gold nanorods with a stealth character for in vivo applications.” Journal of Controlled Release 114, no. 3 (2006): 343–347.

Oh, Seung Wook, Julie A. Harris, Lydia Ng, Brent Winslow, Nicholas Cain, Stefan Mihalas, Quanxin Wang, et al. “A Mesoscale Connectome of the Mouse Brain.” Nature 508, no. 7495 (April 10, 2014): 207–14.

O’Muircheartaigh, J, C Vollmar, G J Barker, V Kumari, M R Symms, P Thompson, J S Duncan, M J Koepp, and M P Richardson. “Focal Structural Changes and Cognitive Dysfunction in Juvenile Myoclonic Epilepsy.” Neurology 76, no. 1 (January 4, 2011): 34–40.

Panyam, Jayanth, and Vinod Labhasetwar. “Biodegradable Nanoparticles for Drug and Gene Delivery to Cells and Tissue.” Advanced Drug Delivery Reviews 55, no. 3 (February 24, 2003): 329–47.

Pardridge, W. M.. “The blood-brain barrier and neurotherapeutics.” NeuroRx: the journal of the American Society for Experimental NeuroTherapeutics 2, no. 1 (2005): 1–2.

Pinzón-Daza, M. L., I. Campia, et al. “Nanoparticle‐ and liposome-carried drugs: new strategies for active targeting and drug delivery across blood-brain barrier.” Curr Drug Metab 14, no. 6 (2013): 625–640.

Rahmim, Arman, Jinyi Qi, and Vesna Sossi. “Resolution Modeling in PET Imaging: Theory, Practice, Benefits, and Pitfalls.” Medical Physics 40, no. 6 (June 2013): 064301.

Reiner, Thomas, Jessica Lacy, Edmund J. Keliher, Katherine S. Yang, Adeeti Ullal, Rainer H. Kohler, Claudio Vinegoni, and Ralph Weissleder. “Imaging Therapeutic PARP Inhibition In Vivo through Bioorthogonally Developed Companion Imaging Agents.” Neoplasia 14, no. 3 (March 2012): 169–177.

Ruggeri, Giacomo, Vera L. Covolan, Marco Bernabò, Li M. Li, Leonardo F. Valadares, Carlos A. P. Leite, and Fernando Galembeck. “Metal Nanostructures with Magnetic and Biodegradable Properties for Medical Applications.” Journal of the Brazilian Chemical Society 24, no. 2 (February 2013): 191–200.

Salata, O.. “Applications of nanoparticles in biology and medicine.” Journal of Nanobiotechnology 2(1) (2004): 3.

Seeman, Philip, Alan Wilson, Peter Gmeiner, and Shitij Kapur. “Dopamine D2 and D3 Receptors in Human Putamen, Caudate Nucleus, and Globus Pallidus.” Synapse (New York, N.Y.) 60, no. 3 (September 1, 2006): 205–11.

Sharma, H. S., S. Hussain, John Schlager, Syed F. Ali, and Aruna Sharma. “Influence of nanoparticles on blood-brain barrier permeability and brain edema formation in rats.” Acta Neurochir Suppl 106 (2010): 359–364.

Shokrollahi, H. “Contrast Agents for MRI.” Materials Science & Engineering. C, Materials for Biological Applications 33, no. 8 (December 1, 2013): 4485–97.

Siegers, G. M., E. J. Ribot, A. Keating, and P. J. Foster. “Extensive expansion of primary human gamma delta T cells generates cytotoxic effector memory cells that can be labeled with Feraheme for cellular MRI.” Cancer Immunol Immunother 62, no. 3(2013): 571–583.

Small, Alex, and Shane Stahlheber. “Fluorophore Localization Algorithms for Super-Resolution Microscopy.” Nature Methods 11, no. 3 (March 2014): 267–79.

Sousa, F., S. Mandal, C. Garrovo, A. Astolfo, A. Bonifacio, D. Latawiec, R. H. Menk, F. Arfelli, S. Huewel, G. Legname, H. J. Galla, S. Krol. “Functionalized gold nanoparticles: a detailed in vivo multimodal microscopic brain distribution study.” Nanoscale 2, no. 12 (2010): 2826–2834.

Srikar, R., A. Upendran, R. Kannan. “Polymeric nanoparticles for molecular imaging.” Wlev Interdiscip Rev Nanomed Nanobiotechnol 6, no. 3 (March 10, 2014): 245–267.

Stoltenburg, R., C. Reinemann, et al. “SELEX-a (Revolutionary method to generate high-affinity nucleic acid ligands.” Biomol Eng. 24, no. 4 (2007): 381–403.

Tam, Jasmine M., Justina O. Tam, Avinash Murthy, Davis R. Ingram, Li Leo Ma, Kort Travis, Keith P. Johnston, and Konstantin V. Sokolov. “Controlled Assembly of Biodegradable Plasmonic Nanoclusters for Near-Infrared Imaging and Therapeutic Applications.” ACS Nano 4, no. 4 (April 27, 2010): 2178–84.

Topalian, S. L., F. S. Hodi, et al. (2012). “Safety, Activity, and Immune Correlates of Anti-PD-1 Antibody in Cancer.” New England Journal of Medicine 366, no. 26: 2443–2454.

Tsien, R. Y., Q. T. Nguyen, M. Whitney (2012). Peptides and aptamers for targeting of neuron or nerves, Google Patents.

Turberfield, A., B. Yurke, and A. P. Mills. “DNA hybridization catalysts and molecular tweezers.” DNA Based Computers \A DIMACS Workshop (Series in Discrete Mathematics and Theoretical Computer Science 54 (2000): 171–182.

Wang, Andrew Z., Vaishali Bagalkot, Christophoros C. Vasilliou, Frank Gu, Frank Alexis, Liangfang Zhang, Mariam Shaikh, et al. “Superparamagnetic Iron Oxide Nanoparticle-Aptamer Bioconjugates for Combined Prostate Cancer Imaging and Therapy.” ChemMedChem 3, no. 9 (September 15, 2008): 1311–15.

Wang, M., J. Etu, and S. Joshi. “Enhanced disruption of the blood brain barrier by intracarotid mannitol injection during transient cerebral hypoperfusion in rabbits.” J Neurosurg Anesthesiol 19, no. 4: 249–56.

Woermann, F. G., S. L. Free, M. J. Koepp, S. M. Sisodiya, and J. S. Duncan. “Abnormal Cerebral Structure in Juvenile Myoclonic Epilepsy Demonstrated with Voxel-Based Analysis of MRI.” Brain: A Journal of Neurology 122, Part 11 (November 1999): 2101–8.

Xu, G., S. Mahajan, et al. “Theranostic quantum dots for crossing blood-brain barrier and providing therapy of HIV-associated encephalopathy.” Front Pharmacol 4 (2013): 140.

Xue, S., J. Qiao, et al. “Design of a novel class of protein-based magnetic resonance imaging contrast agents for the molecular imaging of cancer biomarkers.” Wlev Interdisciplinary Reviews: Nanomedicine and Nanobiotechnology 5, no. 2 (2013): 163–179.

Yamamoto, Seiichi, Hiroshi Watabe, Yasukazu Kanai, Tadashi Watabe, Katsuhiko Kato, and Jun Hatazawa. “Development of an Ultrahigh Resolution Si-PM Based PET System for Small Animals.” Physics in Medicine and Biology 58, no. 21 (November 7, 2013): 7875–88.

Yan, A. C. and M. Levy. “Aptamers and aptamer targeted delivery.” RNA Biol 6, no. 3 (2009): 316–320.

Yurke, B. and A. Mills, Jr‥ “Using DNA to Power Nanostructures.” Genetic Programming and Evolvable Machines 4(2) (2003): 111–122.

Zhang, A., Y. Tu, et al. “Gold nanoclusters as contrast agents for fluorescent and X-ray dual-modality imaging.” J Colloid Interface Sci 372, no. 1 (2012): 239–244.

Zhang, D. Y. and G. Seelig. “Dynamic DNA nanotechnology using strand-displacement reactions.” Nature Chemistry 3, no. 2 (2011): 103–113.

Zhang, D. Y. and E. Winfree. “Control of DNA Strand Displacement Kinetics Using Toehold Exchange.” Journal of the American Chemical Society 131, no. 47 (2009): 17303–17314.

Zhang, Xiao-Dong, Di Wu, Xiu Shen, Pei-Xun Liu, Fei-Yue Fan, and Sai-Jun Fan. “In Vivo Renal Clearance, Biodistribution, Toxicity of Gold Nanoclusters.” Biomaterials 33, no. 18 (June 2012): 4628–38.

Zhang, Xu, Mark R. Servos, and Juewen Liu. “Instantaneous and Quantitative Functionalization of Gold Nanoparticles with Thiolated DNA Using a pH-Assisted and Surfactant-Free Route.” Journal of the American Chemical Society 134, no. 17 (May 2, 2012): 7266–69.

Zhu, Jun, Joshua Chin, Carmen Wängler, Bjoern Wängler, R Bruce Lennox, and Ralf Schirrmacher. “Rapid (18)F-Labeling and Loading of PEGylated Gold Nanoparticles for in Vivo Applications.” Bioconjugate Chemistry 25, no. 5 (May 23, 2014).

